# The long-lasting legacy of reproduction: lifetime reproductive success shapes expected genetic contributions of humans after ten generations

**DOI:** 10.1101/2022.07.26.501566

**Authors:** Euan A. Young, Ellie Chesterton, Virpi Lummaa, Erik Postma, Hannah L. Dugdale

## Abstract

An individual’s lifetime reproductive success (LRS) measures its realised genetic contributions to the next generation, but how well does it predict these over longer periods? Here we use human genealogical data to estimate expected individual genetic contributions (IGC) and quantify the degree to which LRS, relative to other fitness proxies, predicts IGC over longer periods in natural populations. This allows an identification of the life-history stages that are most important in shaping variation in IGC. We use historical genealogical data from two non-isolated local populations in Switzerland to estimate the stabilised IGC for 2,230 individuals ~10 generations after they were born. We find that LRS explains 30% less variation in IGC than the best predictor of IGC, the number of grandoffspring. However, albeit less precise than the number of grandoffspring, we show that LRS does provide an unbiased prediction of IGC and overall predicts IGC better than lifespan and similarly when accounting for offspring survival to adulthood. Overall, our findings demonstrate the value of human genealogy data to evolutionary biology and showing that reproduction - more than lifespan or offspring survival - impacts the long-term genetic contributions of historic humans, even in a population with appreciable migration.

## INTRODUCTION

Fitness is a fundamental concept in evolutionary biology (1). On the individual level, lifetime reproductive success (LRS) - the total number of offspring an individual produces over the course of its lifetime (2)-provides a useful approximation of fitness because it captures the realised ability to contribute genes to the next generation, relative to conspecifics. LRS’s strength stems from, in theory, 1) not confounding selection acting on parents and offspring (3–5) and 2) requiring tracking of only one generation of individuals. LRS is therefore one of the most widely used fitness proxies for the estimation of the strength and direction of natural selection in both free-living and captive populations (6). However, we have little understanding of the extent to which LRS influences the genetic contributions of individuals beyond initial generations (7) and how this compares to other fitness proxies (but see Brommer *et al*. (8)), or in other words, which part(s) of an individual’s life-history are the key determinants of IGC.

An individual’s expected genetic contributions (IGC) – the proportional contribution of an individual to the gene pool at a specific point in time – is expected to stabilise over generations, enabling the estimation of the genetic contributions over far longer periods (7,9,10). Assuming, among others, random mating, non-overlapping generations, negligible inbreeding and stable population size, stabilisation is predicted to occur after approximately 10 generations in a population of 1000 individuals, but longer in larger populations or if any of these assumptions are violated (7,9,11). Largely in line with these theoretical predictions, three studies of wild vertebrate populations found IGC to be relatively stable after around 8 generations (7,10,12) (though other studies have measured expected genetic contributions over shorter periods, e.g. Walle *et al*. (13)). They also found that LRS may predict variation in IGC, but the amount of variation explained varied greatly among studies (<1-48%, Supplementary Table 1) (7,12,14). The latter is expected as an individual’s realised genetic contributions is the ultimate outcome of many factors (e.g. selection, migration, environmental stochasticity and genetic drift), all of which we expect to vary among study systems. For example, we expect LRS to be a poor predictor of IGC if long-term stochastic processes override an initial adaptive response to selection.

The degree to which LRS predicts IGC may also vary with aspects of a species’ or population’s life-history. For example, we could expect the correlation between LRS and IGC to be lower in species – given a similar number of generations – that are long-lived and reproduce over longer time periods, as due to the longer time span there is a greater likelihood that they are exposed to either stochastic mortality events (e.g. a disease outbreak) or changes in selection pressures (e.g. the appearance of a new predator as in Alif *et al*. (12)). However, thus far only species with relatively short generation times (e.g.~2-4 years (7,10,12,14)) have been examined. This is at least partly for practical reasons: estimating IGC and demonstrating their stabilisation is more difficult in longer-lived species because it requires data across greater periods of time.

Human genealogical data, which typically spans centuries rather than decades, provides a powerful opportunity to examine the extent to which LRS predicts long-term genetic contributions in a long-lived species with relatively long generation times. Furthermore, by comparing the predictive power of LRS to other fitness proxies, such as lifespan and the number of grandchildren, we can identify key determinants of variation in IGC. For example, annual survival is considered to be a particularly important driver of within-generation changes to the gene pool (e.g. (15)) in humans. Furthermore, lifespan is associated with increased reproductive success (16). Hence we would expect lifespan to predict IGC, albeit probably with less accuracy than LRS as it does not directly measure reproductive output. The predictive power of LRS is also likely to vary depending on if it is conditioned on offspring survival until a certain age: In pre-demographic transition humans, infant mortality was high (17,18). Therefore, measuring LRS as the number of offspring surviving to adulthood and not only the number born, should better predict IGC. Finally, variation in both the survival and reproduction of an individual and their offspring are ultimately captured by an individual’s number of grandoffspring (19), which is expected to provide a more precise predictor of IGC than LRS. Quantifying the differences in the predictive power of lifespan, number of (surviving) offspring, and the number of grandoffspring will give insight into the relative importance of parental and offspring survival and reproduction in shaping IGC in humans.

The number of grandoffspring is not only expected to explain more variation in IGC (i.e. to be a more precise predictor), but it may also be less biased than LRS (i.e. more accurate). For example, LRS may overestimate IGC if there is an offspring quality versus quantity trade-off or sibling competition, causing offspring from larger families to have lower fitness (20). Conversely, sibling cooperation (e.g. (21) could cause LRS to underestimate IGC if individuals with many siblings have improved fitness. A first step towards identifying the underlying causes of any bias is testing if LRS systematically over- or underestimates IGC. We can do this by quantifying the relationship between an individual’s LRS and the average IGC of their offspring (i.e. of siblings). If this relationship is negative, LRS overestimates the IGC of individuals with high LRS (e.g. due to quality-quantity trade-off or sibling competition), whereas a positive relationship is suggestive of LRS underestimating IGC. This may be the result of e.g. sibling cooperation or parental quality effects (e.g. mediated by socio-economic status) that positively affect both parental reproduction and offspring survival/reproduction (21).

Here, we quantify the degree to which LRS shapes pedigree-derived estimates of stabilised IGC measured after at least 8 generations (10) using data from a genealogical archive containing the life-histories of humans from two parishes in the canton of Glarus, Switzerland. This dataset spans up to 16 generations, containing individuals born in the 16^th^ to the 20^th^ century. We estimate IGC and infer the number of generations required to reach stabilisation. We then use generalised linear mixed models (GLMMs) to examine the degree to which IGCs are predicted by four fitness proxies: lifespan, lifetime reproductive success measured at birth (LRS), LRS counting only offspring surviving to adulthood (LRS_SA_), and the number of grandoffspring. We then compare the predictive power of these four proxies to elucidate the importance of parental and offspring survival and reproduction in shaping IGC, and compare these results to those of previously studied bird species. Finally, we test if LRS provides a biased prediction of IGC by estimating the relationship between an individual’s LRS and the average IGC of their offspring.

## METHODS

### Dataset

We use life-history information, including an individual’s year of birth, marriage and death, and the identity of its children, for individuals born or married in two parishes in the canton of Glarus, Switzerland: Linthal (46°55’N, 9°E) and Elm (46°55’N, 9°10’E). The genealogical archive from which these data were extracted includes records for unmarried adults, children dying before reaching adulthood, and illegitimate children (22) (although these are rare, in line with expectations of historical European populations (23,24)).

The data span over four centuries, containing individuals born from 1562 to 1996. The pedigree reconstructed from these records contained 44,967 individuals, 35,882 maternities, 35,973 paternities and 89,904 full-sibling relationships. The mean maternal and paternal sibship sizes were 4.01 and 4.42, respectively. There were 8,667 founders (individuals with unknown parents), and the mean and maximum pedigree depth was 6.9 and 16 generations, respectively.

During the 18^th^-20^th^ century, population sizes of Linthal and Elm varied between 994-2,645 and 516-1,051, respectively (25,26). The household and family structures are representative of Central Europe as a whole (nuclear and patriarchal), with new households being formed after couples had accumulated enough wealth to get married (27). As such, the median age-at-first reproduction for females was 25, and for 95% of individuals occurred after 19 years of age. For individuals who reproduced, the median number of offspring born was 4 (range = 1-24). Families were largely sustained through the farming of sheep and cattle, with additional earning through weaving and spinning becoming possible in the 18^th^ century (28), particularly in Linthal. Over the course of the entire study period and across all individuals, the median lifespan was 49 years and 74% of individuals lived beyond age 5.

### Estimation of individual genetic contributions

We estimated individual genetic contributions (IGC) following Hunter *et al*. (10), which uses pedigree information to estimate *expected* genetic contributions to future generations, under the *expectation* of random Mendelian segregation of alleles (e.g., each parent contributes 50% of an offspring’s alleles). Hence, IGC provides an estimate of the allele copies given to descendants, and the realised contribution will vary around this expectation. The relatedness matrix, containing the relatedness coefficients between all pairs of individuals (e.g. for a parent and offspring, the relatedness coefficient is 0.5), was created in R 4.1.1 (29) using the package *nadiv* 2.17.1 (30). These relatedness coefficients become expected genetic contributions when directionality is considered: An individual *gives* its offspring 50% of their alleles, and therefore the absolute expected genetic contribution an individual makes to its offspring is 0.5. We will henceforth refer to the individual making the expected genetic contributions as the *focal* individual and to the individual receiving the genetic contribution as the *descendant*.

IGC are equal to the expected genetic contributions proportional to the total gene pool for a given population at a given time point (i.e. all individuals alive and located in the study population). We used birth and marriage locations along with birth and death years to determine if individuals were present in the population (Linthal and Elm were analysed separately) for all individuals with a known birth year (Linthal, N=19558, 98%; Elm, N=16,484, 97%; Supplementary Material 1). To estimate IGC, for each individual in each year, we subset the relationship matrix to include only the focal individual (row) and all individuals present in the specific population at that point (columns), starting at the focal individual’s birth year (or arrival year if an immigrant, see Supplementary Material 1). The total expected genetic contribution of a focal individual to the gene pool in a given year is the sum of this subset of relatedness coefficients. This was done for all the following years until 1990. Following previous studies (7,10,12,14), we did not consider IGC through non-direct descent (e.g., kin genetic contributions) by temporarily removing parental IDs of the focal individuals from the pedigree before creating the relatedness matrix. Genetic contributions were converted into IGC by dividing them by the total number of individuals present in the population in that year.

### Stabilisation of IGC

Although IGCs fluctuate, they are expected to stabilise over time and become representative of longer term genetic contributions (9,11,31). Following previous work (7,12), we evaluated stabilisation of IGC by grouping individuals into 10-year birth cohorts and quantifying the Pearson correlation coefficient between IGC to each subsequent year and the final year considered (1990). 10-year cohorts were used to ensure each cohort had at least 2 focal individuals, as correlation coefficients could not be calculated using one individual When the correlation remained above a 0.95 threshold for a period of 2 generations, IGC were considered to have stabilised. We defined a generation as the mean (± SE) parental age at offspring birth, which were 32.2±0.04 and 32.1±0.05 yr for Linthal and Elm, respectively.

According to this criterion, IGC had stabilised in 1990 for individuals born before 1718 in Linthal (or after 8.5 generations) and before 1734 in Elm (after 8 generations; Figure 1, and see Supplementary Figure 2 for a comparison to non-stabilised IGC). Hence, IGC to the year 1990 from 3475 focal individuals (1,605 from Linthal and 1,870 from Elm) were used for further analyses. The length over which IGC were estimated was at least 274 and 257 yr, and on average 10.1 and 9.9 generations (324.81(±0.86) and 319.26 (±0.93) yr, for Linthal and Elm, respectively), with the birth years of focal individuals ranging between 1575-1734 (Supplementary Figure 3).

**Figure 1:**
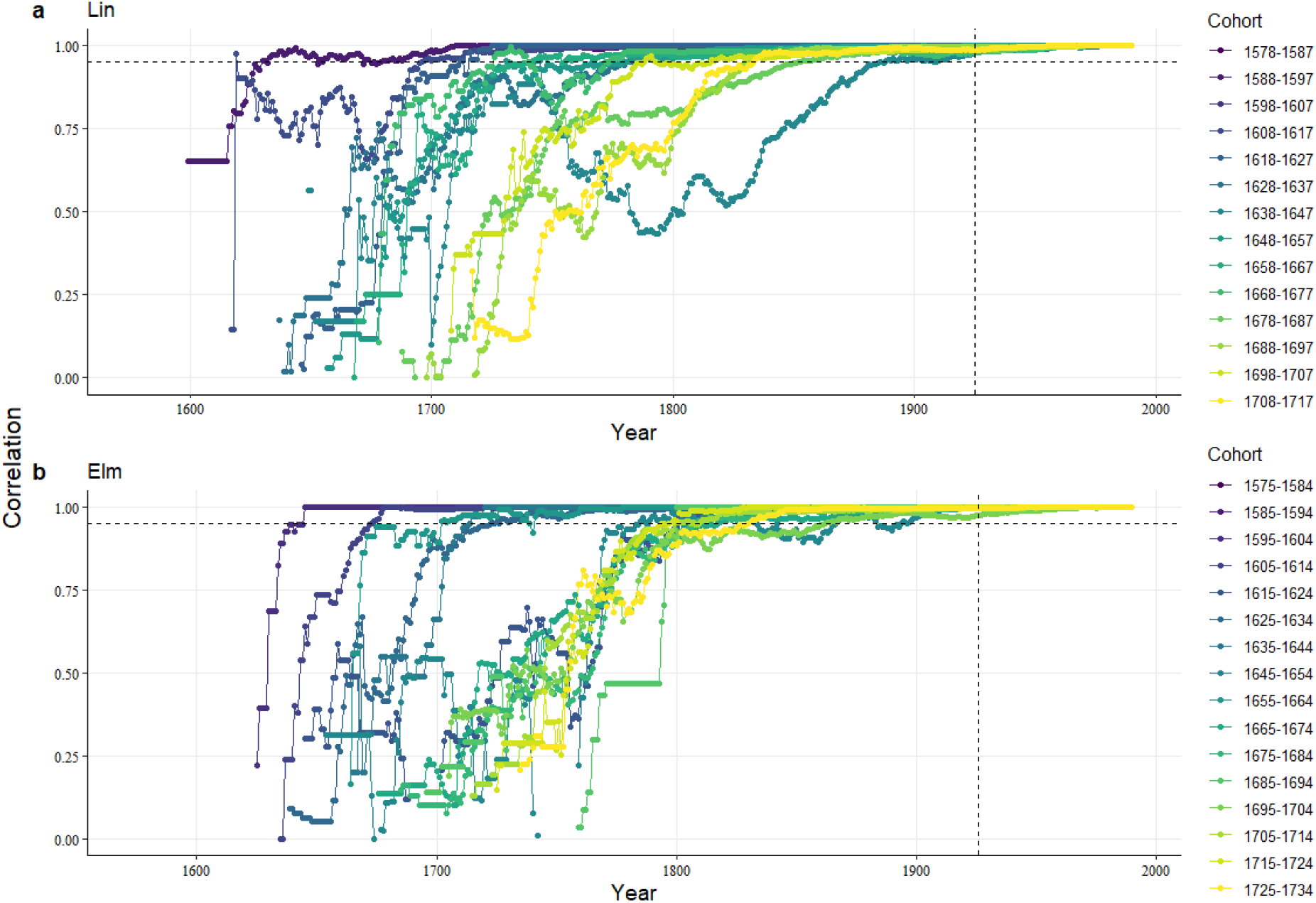
Stabilised IGC. Pearson correlation coefficients between the genetic contribution of individuals grouped into 10-year birth cohort in each year and their final year IGC. Stabilisation is defined as the correlations exceeding 0.95 (horizontal dotted line) for at least two generations pre-1990 (vertical dotted line) (4). Plots are shown for the parishes (**a**) Linthal and (**b**) Elm. Only stabilised cohorts are shown here (born before 1718 for Linthal and 1735 for Elm) but see Supplementary Figure 2.

### Migration

Despite having fulfilled our criterion for stabilisation, IGC will continue to change in populations with a non-zero migration rate (Supplementary Figure 4). This is because immigration decreases IGC by adding to the gene pool but not to the IGC of focal individuals, thereby diluting their contribution to the gene pool. Emigration also decreases IGC and can lead to lineage extinction if emigrating offspring do not contribute to the *local* gene pool. In addition, migration will introduce variation in IGC not captured by any fitness proxies, and hence weakening their correlation with IGC.

To quantify the potential effect of migration on IGC, we classified individuals born and married in the population as residents, individuals born outside but married in the population as immigrants, and individuals born in the population but married outside as emigrants. In Linthal and Elm the vast majority of individuals were residents (62.9% and 61.5%, respectively), but both populations had a substantial proportion of immigrants (16.5% and 15.8%, respectively) and emigrants (20.6% and 22.7%). There was also a very small percentage of individuals who moved between the two parishes (from Linthal to Elm, 0.17%, and Elm to Linthal, 0.22%, see Supplementary Figure 1).

To quantify how often lineage extinction was the result of descendants dispersing versus dying before reproduction, we calculated for each focal individual the percentages of now deceased descendants (traced using the *visPedigree* (32) package) that successfully continued the lineage (i.e. reproduced in the population), did not reproduce in the population, and dispersed (emigrated) out of the population.

### Fitness proxies

We considered the following fitness proxies: lifespan (the difference between the death date and birth date), LRS (lifetime number of offspring produced), LRS_SA_ (lifetime number of offspring surviving to adulthood), and the number of grandoffspring (total number of offspring of an individual’s offspring). Adulthood was defined as the sex-specific 5^th^ percentile of age-at-first reproduction for the whole dataset (females: 19.1 yr, males: 21.2 yr). We estimated lifespan, LRS, and LRS_SA_ for all individuals for which we had an estimated IGC and with known birth and death dates; N=2358), including individuals that died before adulthood. For the number of grandoffspring, we additionally required that the individual’s offspring also had their complete life-history recorded (N=2358).

### Statistical analyses

We used generalised linear mixed models (GLMMs) to examine the relationship between IGC and the four fitness proxies. We used a zero-inflated beta model in which the zero-inflated part of the model modelled the probability of an individual’s IGC to the present-day gene pool being equal to zero (i.e. the probability of lineage extinction) using a logit-link function. The distribution of the non-zero proportional genetic contributions was modelled using a beta distribution.

We controlled for differences in mean IGC, for example due to differences in population size, between both parishes (Linthal or Elm), and the sexes (female or male) by including these as categorical fixed effects. An individual’s 10-year parish-specific birth cohort was fitted as a random intercept to control for temporal variation in mean IGC. We furthermore included a random slope for the effect of each of the fitness proxies to allow their relationship with IGC to vary among parish-specific birth cohorts. Initially a two-way interaction between sex and parish was included, but this was removed if non-significant to aid the interpretation of first-order effects. Model structures were the same for the zero-inflated and beta parts of the model. Counting only individuals that were informative for all predictors, the sample size for these models was 2,230.

To quantify how much variation in IGC each fitness proxy explained, we estimated the Bayesian R-squared for each of our models (33). The significance of the differences in Bayesian R-squared values were evaluated through finding the mode and 95% credible intervals of the difference between the R-squared values of the models being compared (ΔR^2^) and seeing if these 95% credible intervals overlapped 0.

We quantify the bias in LRS in predicting IGC by examining the slope of the relationship between the LRS of an individual and the mean IGC of their offspring. Here we used the same individuals as before, but excluding non-reproducing individuals, leaving 1256 individuals. For this model, we performed a beta regression (with no zero-inflated distribution included) controlling for the same confounding fixed and random effects structures as above. Beta regressions require response variables to non-zero values and we therefore added 10^−10^ to all mean offspring genetic contributions. Here, no relationship would indicate LRS is an unbiased predictor of IGC. We additionally examined if the lifespan of parents was an important covariate, as offspring whose parents died younger might receive less parental care, potentially impacting their IGC.

Both zero-inflated beta and beta models were implemented in the R package *brms* (2.16.1 (34)) using the Markov chain Monte Carlo (MCMC) sampler Rstan (2.21.2 (35)) using R (4.0.2 (29)). For each model, we ran four runs of 6,000 iterations across four cores, sampling every 10 iterations, after a warm-up of 2000 iterations. We set the delta parameter to 0.95 to aid convergence. Default priors were used: flat for all fixed effects and a student’s t distribution for random effects. Convergence of models was confirmed based on R hat parameters and Monte Carlo standard errors being approximately 1 and 0, respectively. The *pp_check* function was used to check that simulated data from the model matched the original data well. We used the probability of Direction (*pd*) (36) (the percentage of the posterior distribution that has the same sign as the median) to infer statistical significance. In line with Makowski *et al*. (36), we classified *pd* values as follows: 0.95-0.975=trend effect; 0.975-0.99 = significant; > 0.99 = highly significant. For random effects, *pd* is not applicable and no significance criteria were used. Figures were created using the packages *brms, ggplot2* (3.3.5, (37)) and *ggpubr* (0.4.0 (38)).

## RESULTS

### Individual genetic contributions (IGC)

We estimated the IGC for 3475 individuals (1,605 from Linthal and 1,870 from Elm), born between 1575-1735, to the individuals making up the gene pool of the parishes of Linthal and Elm in 1990. The probability of an individual’s lineage going extinct was high, with 73% of individuals having an IGC of zero to the 1990 population (Supplementary Figure 5a) (although the extinction rate of specific genes will vary). The majority of extinctions are because an individual did not survive to reproductive age (23.4%), survived until reproductive age but had no offspring (43.5%), had offspring but none survived to adulthood (45.3%), or had surviving offspring but no grandoffspring (52.7%) (Supplementary Figure 5). This leaves approximately 20% of the individual lineage extinctions, which occurred after individuals had at least one grandchild. Over all individuals included in the analysis, a median of 15.9% of their descendants reproduced and thereby continued the lineage, and 14.6% of the descendants had not yet reproduced but were still alive. This leaves 69.7% of the descendants who failed to continue the lineage, of these, a median of 40.3%, due to emigration rather than death without reproducing. Individuals whose lineages did not go extinct on average contributed 0.1% of the genetic material present in the population in 1990 (Supplementary Figure 5a), although one male contributed 0.6% of the Linthal gene pool.

Lifespan, LRS, LRS_SA_, and the number of grandoffspring were positively associated with IGC (Beta distribution, *pd* > 0.975, Table 1, Figure 2). We also found a negative effect of any of the fitness proxies on the probability of an individuals’ lineage going extinct (zero-inflated distribution, *pd* > 0.975, Table 1).

**Table 1:**
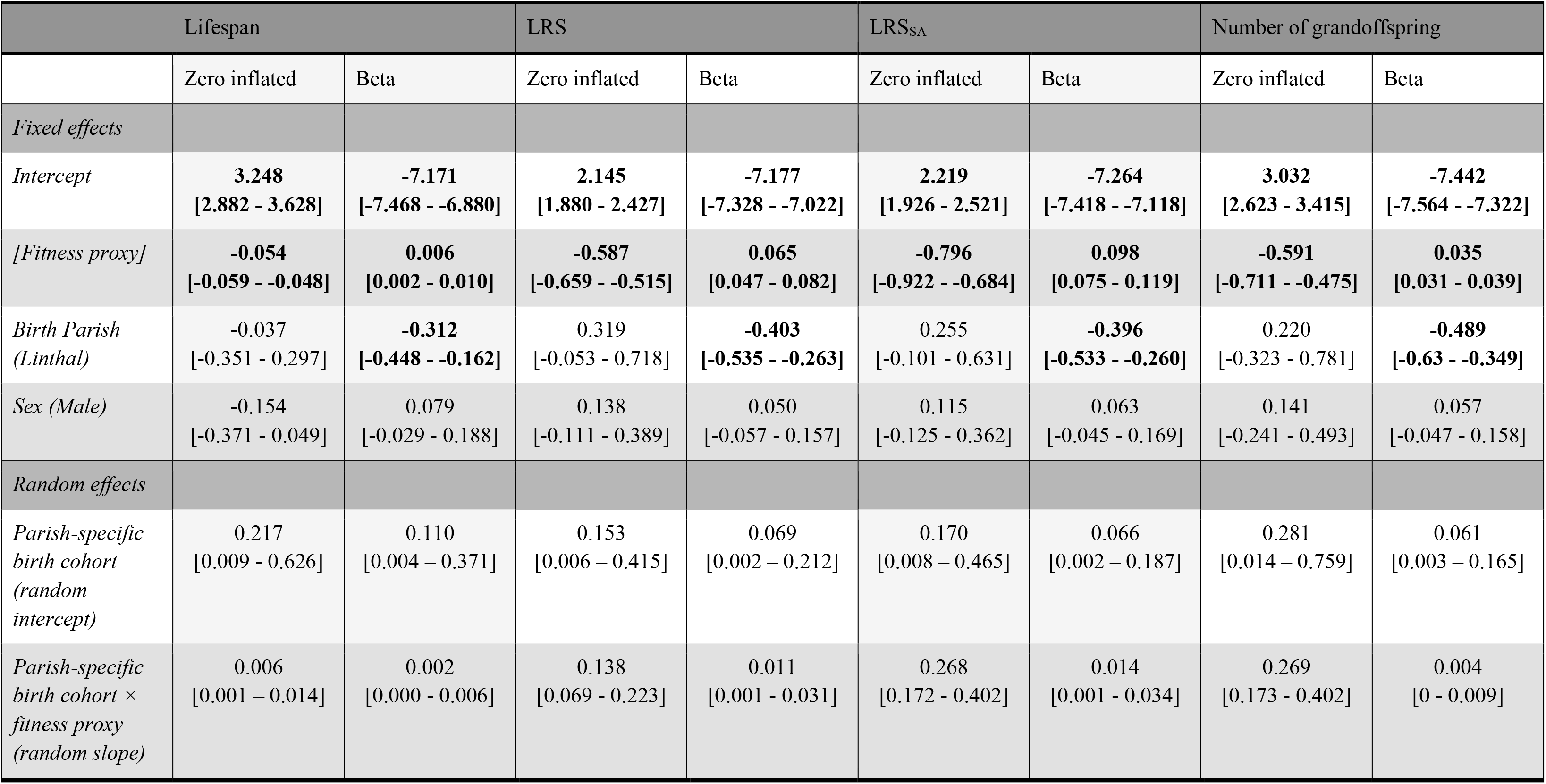
Output from four beta zero-inflated models of IGC with different fitness proxies included: Lifespan, LRS, LRS_SA_, and the number of grandoffspring. Fixed and random effect estimates (posterior distribution median [95% credible intervals] are provided for both the zero-inflated and beta distributions. Significant effects (probability of Direction > 0.975) are in bold. Non-significant two-way interactions were removed from the models. Model results with non-significant interactions included are shown in Supplementary Table 2.

**Figure 2:**
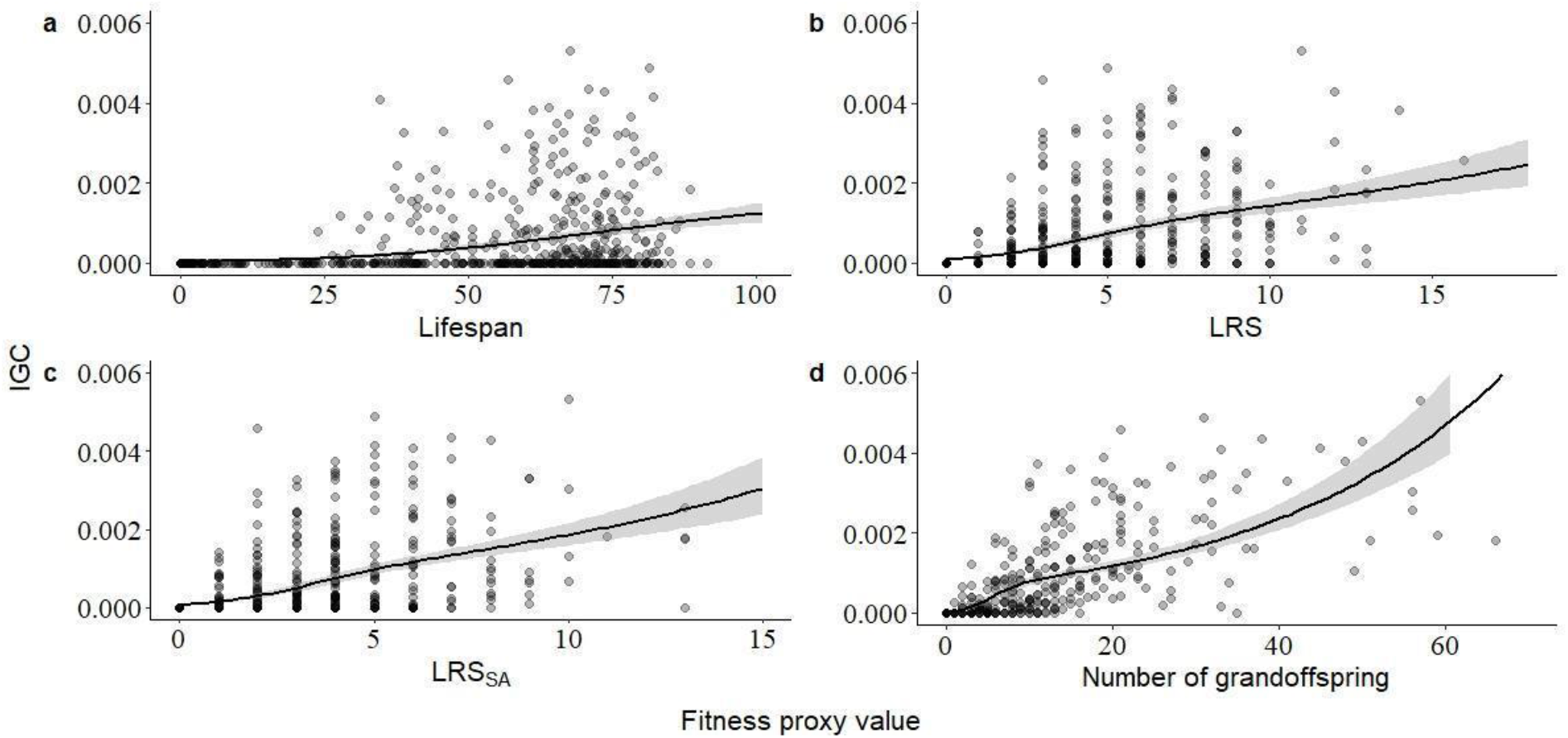
The relationship between IGC and four fitness proxies: (a) lifespan, (b) LRS, (c) LRS_SA_, and (d) the number of grandoffspring. The plots were produced using the *conditional_effects()* function from the R package *brms* to standardise points across values for covariates. Shaded areas indicate 95% credible intervals of the model estimate. Data is conditioned on the mean values of the other predictors (birth cohort, parish and sex). Data points too far away from the values conditioned upon were removed from the plot. Symbols are partially transparent to aid visualisation.

IGC (distribution and extinction probability) were dependent upon several other factors. First, individuals born in Linthal had lower IGC, probably because of its larger population size (all models, beta distribution, *pd* > 0.975, Table 1) and in line with this showed no difference in probability of lineage extinction (zero-inflated distribution, *pd* < 0.975, Table 1). There were no interactions between these effects and sex (*pd* < 0.975, Supplementary Table 2) and no differences between males and females were found (beta and zero-inflated distribution, *pd* < 0.975, Table 1). Further, we found that IGC of individuals varied among birth cohorts (both in their extinction probability and in the non-zero IGC values; see random effects, Table 1). There was also variation among birth cohorts in the slope of the relationship between each fitness proxy and IGC, but except for the slope of the relationship between probability of lineage extinction and LRS, LRS_SA_ and the number of grandoffspring, this variation was small. Finally, a supplementary analysis showed that the proportion of offspring migrating was associated with lower IGC and higher extinction probabilities but this did not substantially change the predictive power of the models (Supplementary Material 2 and Supplementary Table 3).

**Table 2:** On the diagonal, Bayesian R^2^ values (R^2^ and 95% credible intervals) for models containing either lifespan, LRS, LRS_SA_, or the number of grandoffspring and any other significant covariates retained in the model. Pairwise Pearson correlation coefficients (ρ) between fitness proxies are shown above the diagonal (also see Supplementary Figure 6) and the difference in Bayesian R^2^ values are shown below the diagonal (ΔR^2^ and Δ95% credible intervals). Δ95% credible intervals that do not overlap with zero are in bold.

| | Lifespan | LRS | $LRS_{SA}$ | The number of grandoffspring |
| --- | --- | --- | --- | --- |
| Lifespan | $R^2 = 13.2\%$<br>[10.8 - 15.8] | $r = 0.55$ | $r = 0.54$ | $r = 0.43$ |
| LRS | $\Delta R^2 = 14.8\%$<br>**[10.5 - 19.1]** | $R^2 = 27.9\%$ [24.2 - 31.6] | $r = 0.94$ | $r = 0.71$ |
| $LRS_{SA}$ | $\Delta R^2 = 19.2\%$<br>**[14.5 - 23.7]** | $\Delta R^2 = 2.7\%$<br>[-1.8 - 9.2] | $R^2 = 32\%$<br>[27.8 - 36.0] | $r = 0.74$ |
| Number of grandoffspring | $\Delta R^2 = 44.3\%$<br>**[39.5 - 48.2]** | $\Delta R^2 = 29.8\%$<br>**[24.2 - 34.8]** | $\Delta R^2 = 25.2\%$<br>**[19.9 - 31.0]** | $R^2 = 57.3\%$<br>[53.5 - 60.8] |

### How well do fitness proxies predict IGC?

Although all fitness proxies predicted IGC, we found that they significantly varied in their predictive power. As expected, the number of grandoffspring explained most variation in IGC (R^2^=57.3%, Table 2), explaining 44.3 percentage points more variation than lifespan, 29.8 percentage points more than LRS and 25.2 percentage points more than LRS_SA_ (Table 2). Contrary to expectations, the difference in predictability between LRS and LRS_SA_ was very small (ΔR^2^=2.7%, Δ95% Credible Intervals (CrI)=− 1.8% – 9.2%, Table 2). A null model containing no fitness proxy but all other first-order fixed and effects and random effects explained only 1.4% (95% CrI= 0.9% – 2.2%) of the variation in IGC.

### Is LRS an unbiased estimate of IGC?

The per-capita IGC of an individual’s offspring increased with LRS, but the slope of this relationship was very shallow (*pd* > 0.975, posterior mode=0.070, 95% CrI=0.052 - 0.089, Figure 3). This finding suggests that LRS slightly underestimates IGC in larger family sizes. Furthermore, individuals who lived longer had offspring with higher IGC (*pd* > 0.975, posterior mode=0.010, 95% CrI=0.006 - 0.014). As before, and likely due to population size differences, mean IGC of offspring was lower for individuals born in Linthal (*pd* > 0.975, posterior mode= −0.162, 95% CrI=−0.339 - 0.011) but sex differences showed only trend effects and the offspring of males did not have lower mean IGC (*pd* = 0.962, posterior mode=−0.066, 95% CrI=−0.173 - 0.04). No interactions were significant (*pd* < 0.975, Supplementary Table 4).

**Figure 3:**
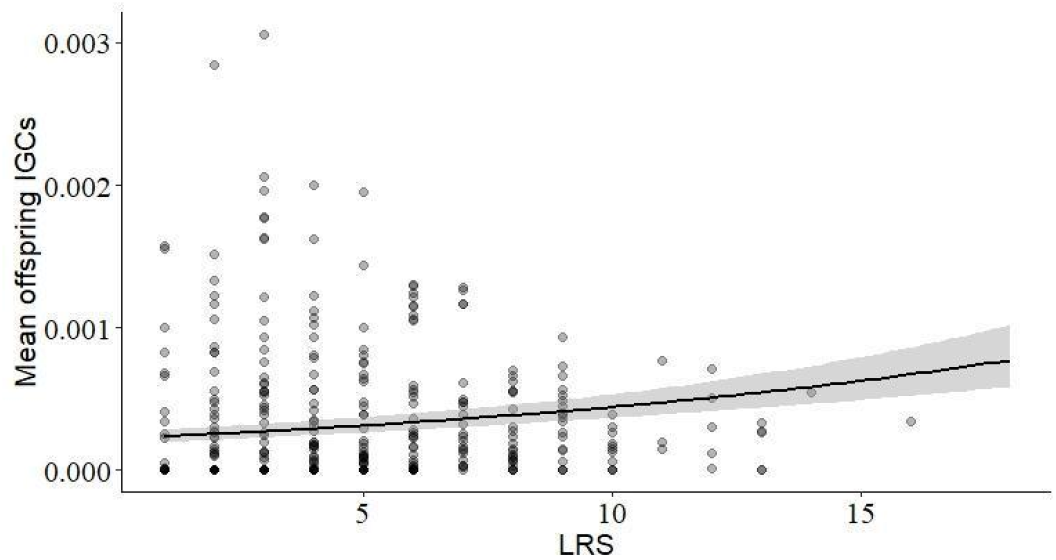
The relationship between mean offspring IGC and LRS. The plots were produced using the *conditional_effects()* function from the R package *brms* to standardise points across values for covariates. Shaded areas indicate 95% credible intervals. Data points too far away from the values specified were removed from the plot and data points included are partially transparent to aid visualisation.

Finally, we found that variance in the mean IGC of offspring was explained by their parents’ birth cohort (posterior mode=0.163, 95% CrI=0.032 - 0.306). The parent’s birth cohort also affected the slope of relationship between LRS and mean offspring IGC but this variation was relatively small (posterior mode= 0.014, 95% CrI=0.001 - 0.038).

## DISCUSSION

We quantified the extent to which LRS and other fitness proxies predict stabilised IGC of individuals measured after ca 10 generations (321 years), in historical humans from the Swiss Canton of Glarus. We found that LRS predicted 28% of the variation in IGC, showing that reproductive success shapes the long-term genetic contributions of individuals even in a population of a long-lived species with appreciable migration that has experienced large and rapid changes in its environment.

We have shown that fitness proxies varied in their predictive power of IGC (Table 2), allowing us to identify the components of an individuals’ life-history that are most important in determining IGC. Overall, the model containing the number of grandoffspring explained 57% of variation in IGC, whereas the next best fitness proxy (LRS_SA_) explained only 32%, followed by LRS (28%) and lifespan (13%). This is broadly in line with results based on genetic contributions estimated over 4 generations in 19^th^ century Sweden (39). That the number of grandoffspring explained the most variation was expected, as the number of grandoffspring incorporates the most information about the life-history of an individual. However, together with our finding that LRS_SA_ and LRS explain a similar amount of variation in IGC (28% vs 32%) and that lifespan explains only 13% of the variation in IGC, this suggests that offspring mating and reproduction is a much greater determinant of IGC than survival (of both offspring and the individual themselves), even in a population with substantial childhood mortality (Supplementary Figure 1). Finally, we showed no difference between the sexes although the highest IGCs were multiple-marrying males who’s first wives died around the age of menopause, allowing the widowers to remarry a younger female and achieve a lifetime reproductive success (and IGC) greater than males who did not remarry.

Although the number of grandoffspring explains the most variation in IGC, the number of grandoffspring is not necessarily the most useful fitness proxy. First, although statistically significant, we found a weak relationship between LRS and the average IGC of their offspring, showing that LRS is a relatively unbiased measure of IGC. The positive association suggests that the increase in predictive power between LRS and number of grandoffspring is not due to LRS being a biased predictor. The number of grandoffspring will naturally be a more precise predictor of IGC than LRS because it is closer in time to IGC and therefore incorporates more of the stochasticity that influences IGC. Albeit small, the positive relationship between LRS and per-capita IGC argues against the existence of an offspring quality-quantity trade off, which has been previously found in humans (40–42). Instead it is somewhat suggestive of positive sibling effects, perhaps due to alloparenting (21), or an overriding effect of parental quality (e.g. socio-economic status) (e.g. (43,44)). Second, there are practical reasons that limit the utility of the number of grandoffspring as a fitness proxy: Not only is it more sampling intensive, reliably counting the number of grandoffspring may not be feasible if a significant proportion of the population disperses outside of the study site, or offspring cannot be linked to parents once they have reached independence. Third, the number of grandoffspring confounds the fitness of multiple individuals, which can be problematic when estimating the strength of phenotypic selection (3–5). All things considered, our results therefore strengthen the case for LRS as an evolutionary relevant and relatively unbiased fitness proxy when it comes to the study of selection in humans, assuming our findings are representative for other populations and time periods.

Although at first sight high, our finding that 70% of individual lineages went extinct over the study period is similar to that found in previous studies on pedigreed populations of birds, which reported extinction probabilities of 61-71% (Supplementary Table 1), and comparable with levels of lineage extinction in bighorn sheep (13) in humans after four generations in Sweden (45). The main difference between our results and those for the three bird studies (7,12,14) was that when we measured LRS at a later point in the offspring’s life (i.e. LRS_SA_) the ability of LRS to predict IGC did not increase greatly. This is in contrast to, for example (14), which found that offspring survival was a key determinant of reproductive success. Our results therefore suggest that despite substantial infant mortality, offspring survival to adulthood is a less important determinant of IGC than mating and reproductive success in humans.

The amount of variation in IGC explained by LRS measured close to birth (28%) in this study was close to previous findings for song sparrows and scrub-jays (37% and 32%, respectively), but higher than house sparrows (0-4%). This is somewhat surprising given the likely negligible role of migration in the latter island population but is potentially due to a bottleneck in the population that occurred between fitness proxies being measured and the estimation of IGC (12). This could have caused stochastic mortality resulting in low predictive power of fitness proxies. Another likely factor explaining different findings across populations is the role of stochasticity in driving variation in LRS itself. LRS is influenced by both environmental and genetic components with the environment contributing most of the variation (46), including in humans (47). In species where the environment determines less variation in LRS, LRS would be expected to be a greater predictor of IGC (7,8). Here, we showed that environmental effects were an important factor, with non-negligible variation in IGCs being explained by an individual’s birth cohort (Table 1). Further, as mean LRS values decrease there is a greater likelihood of lineages going extinct due to stochasticity, drift or dispersal (7,48), which perhaps partially explains the relatively high rates of lineage extinction in this study (70% vs 61-71%, Supplementary Table 1). Future studies could examine if this phenomenon is detectable across the human fertility transition towards lower LRS. In summary, there are both similarities and differences across study systems, but the small number of species and the lack of different human populations (across cultures) studied limits broader extrapolation.

72% of the variation in IGC remained unexplained in the model containing LRS, with migration being a contributing factor to this unexplained variation: First, there were significant levels of both immigration and emigration (Supplementary Figure 1), and both are expected to decouple the relationship between LRS and IGC. Dispersal of descendants of ancestral individuals is a particularly important driver of lineage extinction. Although migration is also expected to reduce the stabilisation times relative to theoretical expectations, we observed stabilisation times lower than theory predicts (9). One explanation for this is that the effective population size (number of breeding individuals) is far lower than the total population size, for example because a significant proportion of individuals didn’t reproduce (see Supplementary Figures 1 and 5). However, other explanations (e.g. non-random mating) are also possible and it is clear that we need to further our understanding of the drivers of the stabilisation of IGC in natural populations. Finally, the slopes of the relationships between fitness proxies are generally low (0.006 to 0.098); thus, discrepancies between LRS and fitness proxies generally arose because of individuals having high LRS but low or no IGC, and not vice-versa. Here, migration is a likely driver, although other evolutionary forces will be at play (e.g. drift and fluctuating selection). Enumerating the relative contributions of these factors across different systems (or using simulations, e.g. (49)) should be a target of future work.

Although still in its infancy, the use of pedigree data to estimate long-term genetic contributions opens a range of exciting avenues. Building on our work using human genealogical data, and the work on non-human animals by others (7,12,14), future work would benefit from further exploration of the similarities and differences among the different methodologies at our disposal. In particular between gene dropping methods (7,12,14) and expected genetic contributions (e.g. (10), this study), as the two will not necessarily equate. Furthermore, while our study has highlighted the ability of human genealogical data to provide insight into human evolution (50,51), and the estimation of fitness more broadly, applying these methods to similar data for an array of human populations (see (52), for a review) will allow us to quantify the degree to which these findings hold across cultures, environments and time.

## COMPETING INTERESTS

Nothing to declare

## FUNDING

E.Y.’s PhD was funded by the University of Groningen, through a Rosalind Franklin Fellowship awarded to H.L.D. Digitisation and transcription of the data were funded by the Swiss National Science Foundation (31003A_159462). VL was funded by the Strategic Research Council of the Academy of Finland (grant number 345185 and 345183).

## ACKNOWLEDGEMENTS

We thank Beat Mahler and Fritz Rigendinger of the Landesarchiv des Kantons Glarus for enabling access to the data; Darren Hunter for providing the initial code for estimating IGC; and Aïda Nitsch, Milla Salonen, and Simon Evans for insightful comments on the manuscript. Finally, we thank the editor and two anonymous reviewers for constructive feedback that considerably improved the manuscript.

## Notes

### Competing Interest Statement

The authors have declared no competing interest.

### Summary of Updates

This version of the manuscript has been revised following reviewer comments. Statistical models and results have remained approximately the same but the interpretation and general framing of the results have been modified.

