## Supplementary material for "The long-lasting legacy of reproduction: lifetime reproductive success shapes expected genetic contributions of humans after ten generations"

**Supplementary Table 1: Summary of previous estimates of the variance in IGC explained by different fitness proxies (measured as R-squared)**. Unless specified otherwise, fitness proxies were defined as follows: Lifespan, difference between death year and birth year of individual; LRS (number of eggs), individual lifetime number of eggs produced; LRS (hatchlings), individual lifetime number of offspring produced counted after 2 days; LRS (ringed), individual lifetime number of offspring produced counted when chicks were ringed (note that ringing date varied between studies); LRS (independent), individual lifetime number of offspring produced counted 24 days post hatch; LRS (recruited) LRS (hatchlings), individual lifetime number of offspring produced counted after 1 year; and number of (great) grandoffspring, total number of (great) grandoffspring of an individual. The sample sizes are recorded as the number of individuals used in analysis. The study period was the length between measurement of fitness proxies and IGC. Where studies reported sexes separately, males are specified in parentheses. Study period is defined as the number of years between the measurement of an approximation of an individual's fitness and the estimation of IGC. The extinction probability is measured as. the percentage of individuals that reported zero IGC. For Reid *et al.* (2019), adjusted R-squared values are reported. For Alif *et al.* (2021), values were approximated from Figure 2 in paper. Cells where data were not reported are left blank.

| Study species | Song sparrows (*Melospiza melodia*) | Floridian Scrub-Jay (*Aphelocoma corulescens*) | House Sparrow (*Passer domesticus*) | Humans from the Swiss canton of Glarus |
|---|---|---|---|---|
| Reference | Reid *et al.* (2019) | Chen *et al.* (2019) | Alif *et al.* (2022) | This paper |
| Sample size | 55 (94) | 926 | 86 | 2,230 |
| Study period | 20 years | 14-26 years | 16-20 years | 257-415 years |
| Extinction probability | 67% (71%) | | 61% | 73% |
| *Fitness proxy* | *R-squared* | | | |
| Lifespan | 0.25 (0.29) | | | 0.132 [0.11 – 0.16] |
| LRS (number of eggs) | | | 0.00025† | |
| LRS (offspring born) | | | | 0.28 [0.24 – 0.32] |
| LRS (hatchlings) | | | 0.00027† | |
| LRS (ringed) | 0.37 (0.30)‡ | 0.32†§ | 0.04†¶ | |
| LRS (independent) | 0.36 (0.31) | | | |
| $LRS_{SA}$ | | | | 0.32 [0.28 – 0.36] |

| | | | | |
|---|---|---|---|---|
| LRS (recruited) | 0.48 (0.55) | | 0.21†¶¶ | |
| The number of grandoffspring | | 0.69† | | 0.57 [0.54 – 0.61] |
| Number of great-grandoffspring | | 0.76† | | |

†R-squared approximated by squaring the reported correlation coefficients.

‡Ringing occurred in 6 day old nestlings.

§Ringing occurred in 11 day old nestlings.

¶Ringing occurred in 12 day old nestlings.

¶¶One year olds that bred within the population.

¶¶¶Bayesian R-squared values [95% credible intervals]

SUPPLEMENTARY MATERIAL 1

Estimating IGC requires knowledge of when each individual is alive and present (i.e. located) in the population, and is thereby part of the population's gene pool, at each time point. For this we used recorded birth and marriage locations of individuals. Birth and marriage locations are known for most individuals (Linthal: birth location = 17,351/19,558, marriage location = 10,132; Elm: birth location = 14,787/16,484, marriage location = 8,571), including those that married in other parishes within the Canton of Glarus. For individuals without a recorded death year (Linthal = 7,868; Elm, N = 7,547), the sex-specific median lifespan for the individual's 10-year birth cohort was added to their birth year to estimate their death year. Median lifespans for the 10-year cohorts decreased after 1920, due to a sampling bias in recorded lifespan towards individuals dying at younger ages. Therefore, individuals born after 1920 with missing death years had their death year estimated based on the 1910-1920 sex-specific median lifespan (Linthal, N=3,115; Elm, N=2,843).

There is appreciable migration in the populations (Supplementary Figure 1), so we classed individuals into different 'migration statuses' and estimated their arrival and departure years according to the following criteria. Firstly, individuals who had recorded birth and marriage locations in that parish (*residents married*), arrival and departure years for individuals were assumed to be equal to their birth and death years (Linthal = 5,644; Elm = 4,877). This was also done for individuals dying before reaching adulthood (defined by the 20-year cohort sex-specific $5^{th}$ percentile of age-at-first reproduction and ranged from 18.7-23.9 years) (Linthal = 4,046; Elm = 2,631). Finally, individuals that were expected to still be alive (i.e. born within one median sex-specific lifespan for 1910-20 of 1990; Linthal=1320, Elm= 1618) and either had a recorded marriage location in the particular population or were born after 1946 (after which records of marriage decrease) were assigned a departure year as their death year (when available) or as 1991. Thereby effectively being treated as still alive in further analyses.

Individuals with a recorded birth in the respective population but with a recorded marriage in another parish in Glarus and a recorded death year (*emigrants*) were assigned arrival years as their birth years and departure years as the birth year of their first offspring born outside the parish. For descriptive purposes we separated *emigrants* who emigrated to one of the other study populations (Linthal and Elm) and *emigrants* who emigrated to one of the other parishes in Glarus. Individuals born in the study population but that had no marriage or death record – which is likely to arise if they have emigrated outwith Glarus – were additionally classed separately and their departure years were predicted by adding their birth recorded birth year to the median age-at-first reproduction for their particular 20-year birth cohort. Thus, the emigrant classifications were for Linthal: *emigrants (to Elm)* (N=33), *emigrant (to other Glarus parish)* (N=1217), and *emigrant (to outside Glarus)* (N=2799); and for Elm were *emigrant (to Linthal)* (N=37), *emigrant (to other Glarus parish)* (N=1056), and *emigrant (to outside Glarus)* (N=2669).

In a similar manner, individuals with a missing recorded birth location outwith the respective population but with a recorded marriage in the population and a recorded death year (*immigrants*) were assigned arrival years as the birth year of their first offspring within the parish and departure years equalled their recorded death years. Again, we separated *immigrants* who had recorded births in one of the other study populations and other parishes in Glarus. Individuals with no recorded birth record location but a marriage record within the population in question were assumed to have been born outside Glarus as it would be unlikely for an individual who had married within a particular population to have also been born there but this not to have been recorded. Thus, the immigrant classifications were: for Linthal *immigrants (from Elm)* (N=41), *immigrant (from other Glarus parish)* (N=990), and *immigrant (from outside Glarus)* (N=2207); and for Elm were *immigrant (from Linthal)* (N=35), *immigrant (from other Glarus parish)* (N=878), and *immigrant (from outside Glarus)* (N=1698).

Any individual that had a recorded birth year and location in the particular population and had a recorded death year but no marriage record was assumed to have stayed in the population but remained unmarried (*resident (unmarried)*, Linthal=1261, Elm=995). Finally, there were also a small proportion individuals who had married multiple times in Linthal and Elm (N=560 and N=452, respectively). In all cases these individuals were born in the study parish but married and reproduced first outside of the population in question and thus for this period these individuals were classed as *emigrants* in accordance with the criteria above. If these individuals then re-entered the population they were born in they were assigned a new migrant class (*resident returned* Linthal=85 and Elm=62) and their arrival year was estimated as the first birth year of their offspring within the parish they were born in and departure years as their death year. The total number of individuals belonging to each migration class in each parish are shown in Supplementary Figure 1.

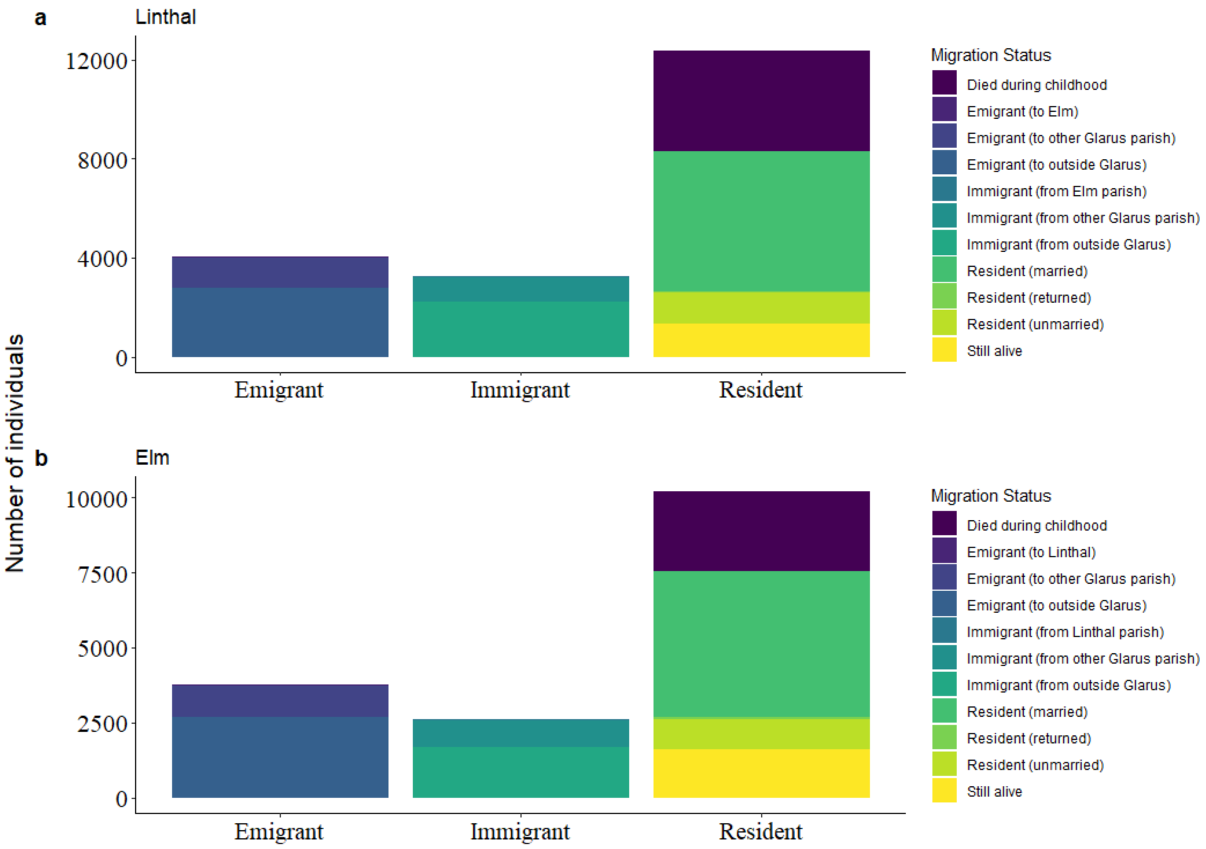

**Supplementary Figure 1: Barplot showing the number of individuals of each migration status in a) Linthal and b) Elm.** Migration status was assigned according to the criteria in Supplementary Material 1. These follow three broad categories (x-axis, *emigrant*, *immigrant*, *resident)* and 11 subcategories (colours, see legend).

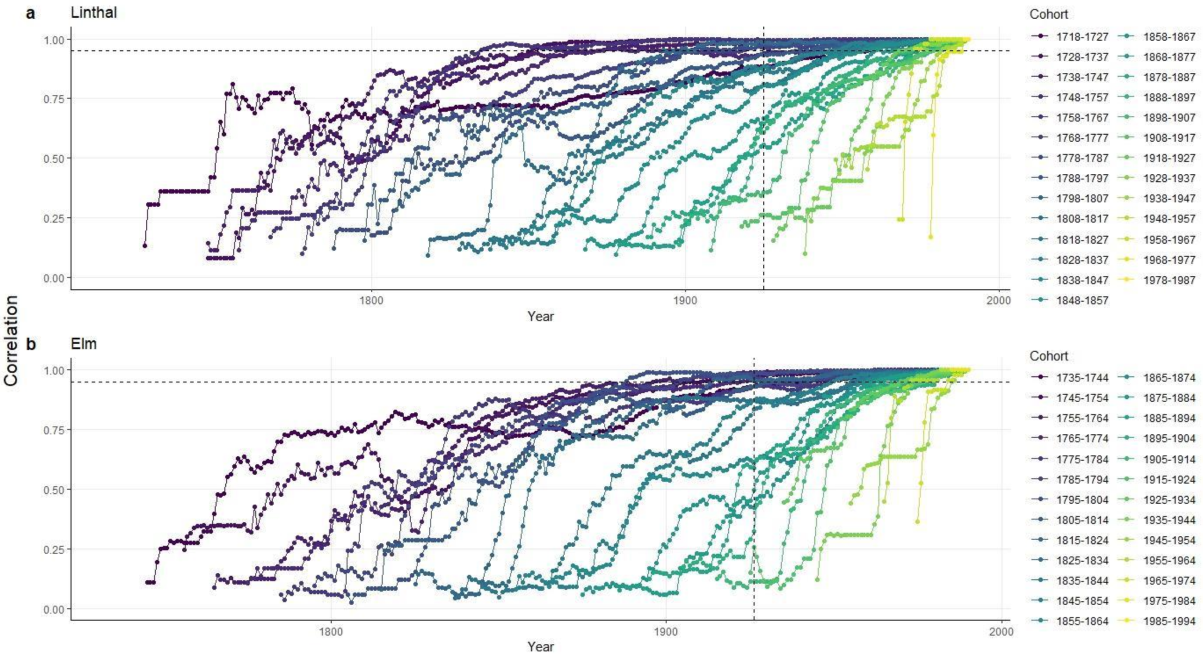

**Supplementary Figure 2: Non-stabilised IGC.** Pearson correlation coefficient between the genetic contribution of each 10-year birth cohort to each subsequent cohort and its genetic contribution to the final cohort. Stabilisation is defined as the correlations exceeding 0.95 (horizontal dotted line) for at least two generations pre-1990 (vertical dotted line) (4). Plots are shown for the parishes (**a**) Linthal and (**b**) Elm. Only cohorts after 1717 for Linthal and 1734 for Elm are shown.

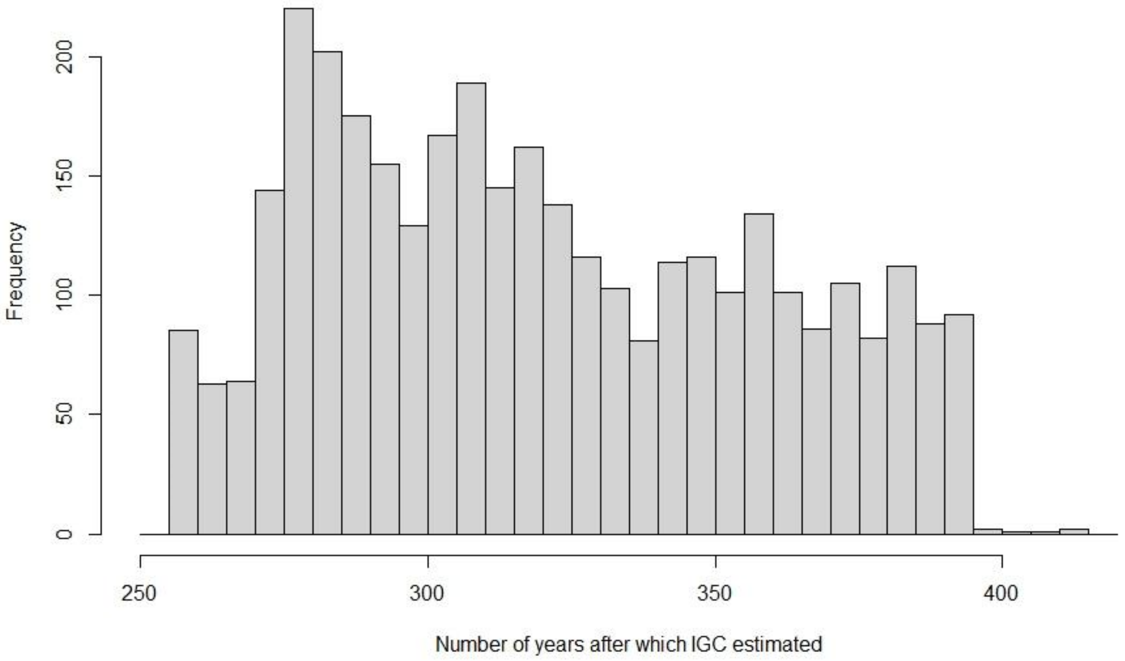

**Supplementary Figure 3: Histogram showing the number of years over which IGC were estimated.** Calculated as the year estimated in (i.e. 1990), minus the individual's birth or arrival year.

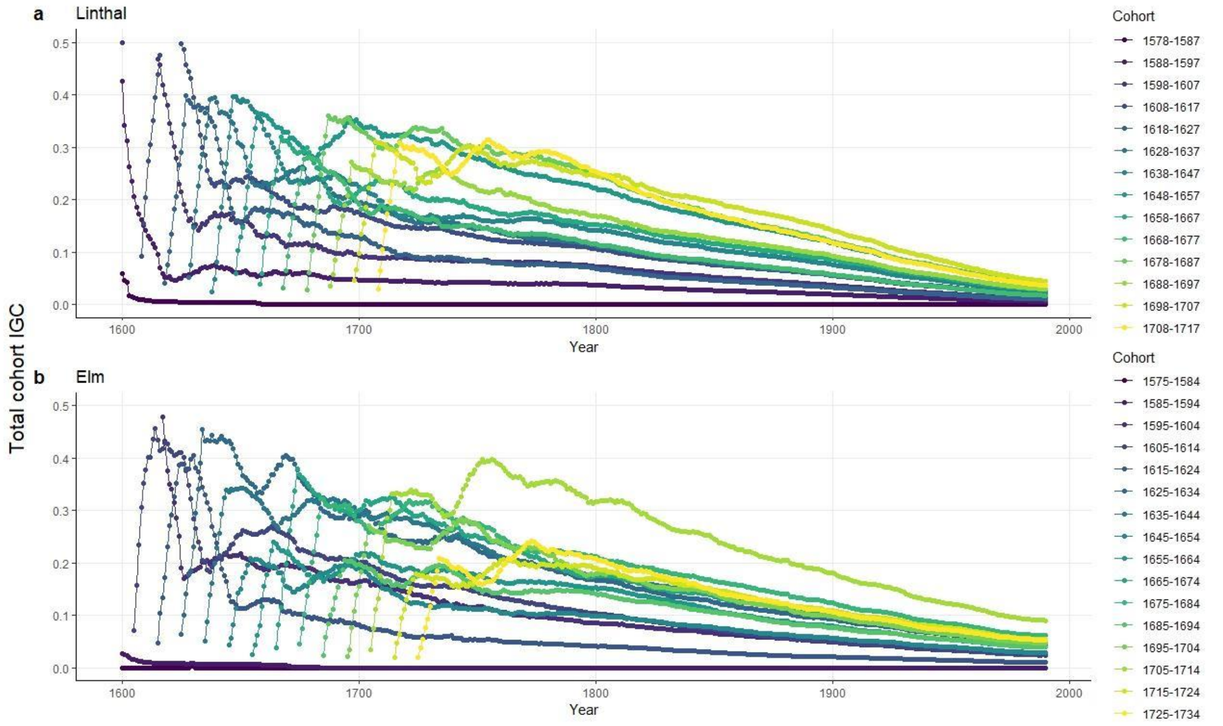

**Supplementary Figure 4: Cohort IGC over time.** Total 10-year cohort IGC for (**a**) Linthal and (**b**) Elm from 1600-1990. Only cohorts used in the analyses are shown.

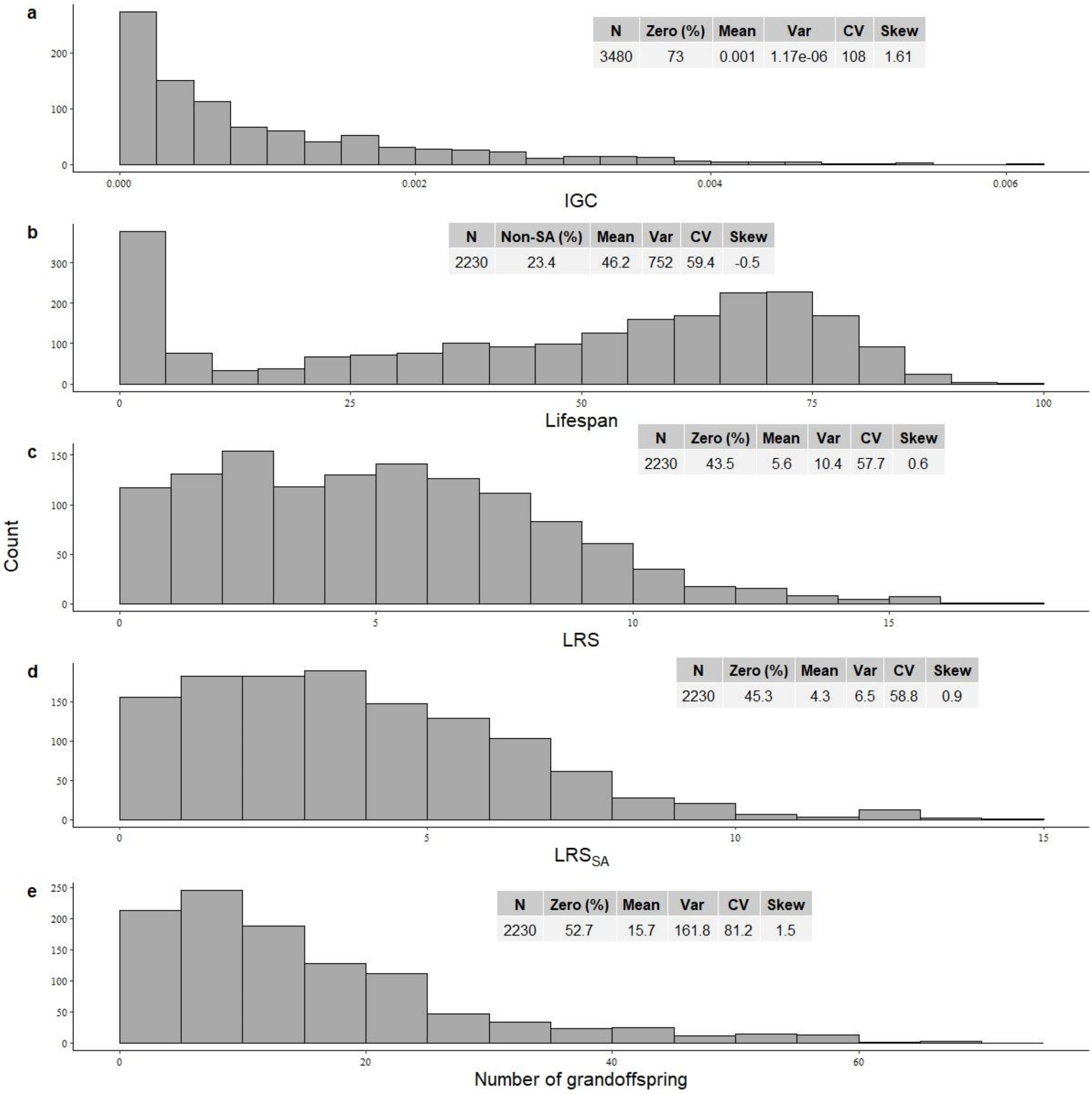

**Supplementary Figure 5: Distribution of IGC and fitness proxies.** Histograms showing the distribution of (**a**) individual genetic contributions (IGC) and fitness proxies: (**b**) lifespan, (**c**) LRS, (**d**) $LRS_{SA}$, and (**e**) the number of grandoffspring. Zero values were removed from plots to aid visualisation. Boxes display several descriptive statistics for the data: N, the total sample size; Zero (%), percentage zeros; Mean, mean value; Var, raw variance, CV, coefficient of variance (calculated as standard deviation/mean multiplied by 100); Skew, the distribution skewness. The Mean, Var, CV, and Skew describe only the non-zero data. For lifespan, the percentage of individuals dying before adulthood (Females=19.1, Males=21.2 yr) are shown, instead of the percentage of zeros.

**Supplementary Table 2: Output from the four beta zero-inflated models of IGC with one of the following fitness proxies included: Lifespan, LRS, $LRS_{SA}$, or the number of grandoffspring, including non-significant interaction terms.** Reported are the fixed and random effect estimates (posterior distribution median), their 95% credible intervals (95% CrI); and the probability of Direction (*pd*) (not available for random effects). The output from both the zero-inflated and beta distributions are shown. Significant effects (*pd* > 0.975) are shown in bold.

| | Lifespan | | LRS | | $LRS_{SA}$ | | Number of grandoffspring | |
|---|---|---|---|---|---|---|---|---|
| | Zero inflated | Beta | Zero inflated | Beta | Zero inflated | Beta | Zero inflated | Beta |
| Fixed effects | | | | | | | | |
| Intercept | **3.22 [2.845 - 3.618]** | **-7.152 [-7.451 - -6.859]** | **2.113 [1.823 - 2.417]** | **-7.161 [-7.314 - -6.995]** | **2.182 [1.871 - 2.486]** | **-7.251 [-7.402 - -7.099]** | **3.012 [2.588 - 3.438]** | **-7.435 [-7.566 - -7.303]** |
| [Fitness proxy] | **-0.054 [-0.059 - -0.048]** | **0.006 [0.001 - 0.01]** | **-0.589 [-0.661 - -0.519]** | **0.065 [0.046 - 0.083]** | **-0.798 [-0.922 - -0.675]** | **0.098 [0.077 - 0.12]** | **-0.593 [-0.712 - -0.481]** | **0.035 [0.031 - 0.039]** |
| Birth Parish (Linthal) | 0.021 [-0.368 - 0.46] | **-0.342 [-0.539 - -0.145]** | 0.415 [-0.04 - 0.884] | **-0.44 [-0.625 - -0.263]** | 0.342 [-0.112 - 0.821] | **-0.428 [-0.611 - -0.24]** | 0.305 [-0.362 - 0.989] | **-0.497 [-0.663 - -0.318]** |
| Sex (Male) | -0.115 [-0.389 - 0.144] | 0.059 [-0.083 - 0.205] | 0.214 [-0.089 - 0.518] | 0.024 [-0.114 - 0.163] | 0.196 [-0.119 - 0.5] | 0.044 [-0.097 - 0.183] | 0.204 [-0.252 - 0.678] | 0.048 [-0.08 - 0.174] |
| *Parish (Linthal) × Sex (Male)* | -0.094 [-0.487 - 0.321] | 0.047 [-0.19 - 0.274] | -0.193 [-0.702 - 0.302] | 0.065 [-0.158 - 0.289] | -0.187 [-0.703 - 0.347] | 0.052 [-0.167 - 0.277] | -0.178 [-0.938 - 0.6] | 0.019 [-0.174 - 0.224] |
| Random effects | | | | | | | | |
| Parish-specific birth cohort (random intercept) | 0.227 [0.007 - 0.655] | 0.104 [0.005 - 0.348] | 0.156 [0.006 - 0.451] | 0.07 [0.003 - 0.218] | 0.17 [0.007 - 0.449] | 0.068 [0.003 - 0.187] | 0.284 [0.014 - 0.75] | 0.061 [0.003 - 0.169] |
| Parish-specific birth cohort × fitness proxy (random slope) | 0.006 [0.001 - 0.014] | 0.002 [0 - 0.006] | 0.139 [0.065 - 0.22] | 0.011 [0 - 0.03] | 0.267 [0.165 - 0.393] | 0.014 [0.001 - 0.036] | 0.268 [0.174 - 0.4] | 0.004 [0 - 0.009] |

SUPPLEMENTARY MATERIAL 2

We carried out a supplementary analysis of the impact of offspring dispersal to check if this changed our results. Here, we additionally controlled for the proportion of an individual's offspring which did not disperse in our models (calculated as the number of an individual's offspring who remained in their birth parish ÷ the total number of offspring). Although individuals who had a greater proportion of non-dispersing offspring had higher extinction probabilities and lower IGC, these did not change the results qualitatively (Supplementary Table 3).

**Supplementary Table 3: Output from the four beta zero-inflated models of IGC with one of the following fitness proxies included: Lifespan, LRS, $LRS_{SA}$, or the number of grandoffspring, including non-significant interaction terms.** Reported are the fixed and random effect estimates (posterior distribution median), their 95% credible intervals (95% CrI); and the probability of Direction (*pd*) (not available for random effects). The output from both the zero-inflated and beta distributions are shown. Significant effects (*pd* > 0.975) are shown in bold.

| | Lifespan | | LRS | | $LRS_{SA}$ | | Number of grandoffspring | |
|---|---|---|---|---|---|---|---|---|
| | Zero inflated | Beta | Zero inflated | Beta | Zero inflated | Beta | Zero inflated | Beta |
| Fixed effects | | | | | | | | |
| Intercept | **1.911 [1.136 - 2.662]** | **-7.83 [-8.279 - -7.417]** | **2.317 [1.566 - 3.072]** | **-8.066 [-8.469 - -7.672]** | **3.317 [2.452 - 4.195]** | **-8.442 [-8.862 - -8.039]** | **1.231 [0.327 - 2.164]** | **-7.924 [-8.272 - -7.588]** |
| [Fitness proxy] | **-0.052 [-0.059 - -0.047]** | **0.005 [0.001 - 0.009]** | **-0.590 [-0.663 - -0.522]** | **0.069 [0.051 - 0.086]** | **-0.825 [-0.953 - -0.702]** | **0.111 [0.09 - 0.133]** | **-0.583 [-0.704 - -0.474]** | **0.035 [0.031 - 0.039]** |
| Birth Parish (Linthal) | -0.010 [-0.337 - 0.358] | **-0.307 [-0.448 - -0.162]** | 0.323 [-0.037 - 0.709] | **-0.398 [-0.524 - -0.264]** | 0.272 [-0.099 - 0.642] | **-0.391 [-0.526 - -0.264]** | 0.169 [-0.319 - 0.723] | **-0.479 [-0.614 - -0.339]** |
| Sex (Male) | -0.151 [-0.360 - 0.049] | 0.073 [-0.036 - 0.177] | 0.130 [-0.098 - 0.369] | 0.046 [-0.060 - 0.154] | 0.123 [-0.125 - 0.381] | 0.059 [-0.045 - 0.171] | 0.135 [-0.227 - 0.490] | 0.05 [-0.051 - 0.152] |
| *Proportion of non-migrating offspring* | **1.361 [0.711 - 2.063]** | **0.769 [0.423 - 1.141]** | -0.177 [-0.914 - 0.54] | **0.947 [0.547 - 1.348]** | **-1.128 [-1.968 - -0.279]** | **1.212 [0.816 - 1.623]** | **1.993 [1.129 - 2.929]** | **0.532 [0.190 - 0.893]** |
| Random effects | | | | | | | | |
| Parish-specific birth cohort (random intercept) | 0.215 [0.007 - 0.625] | 0.103 [0.003 - 0.346] | 0.149 [0.006 - 0.412] | 0.070 [0.002 - 0.212] | 0.169 [0.007 - 0.463] | 0.061 [0.002 - 0.176] | 0.298 [0.010 - 0.772] | 0.066 [0.004 - 0.171] |
| Parish-specific birth cohort × fitness | 0.006 [0.001 - 0.014] | 0.002 [0.000 - 0.005] | 0.139 [0.069 - 0.225] | 0.010 [0 - 0.030] | 0.272 [0.172 - 0.405] | 0.01 [0 - 0.028] | 0.261 [0.170 - 0.395] | 0.005 [0 - 0.010] |

| |
|---|
| proxy (random slope) |

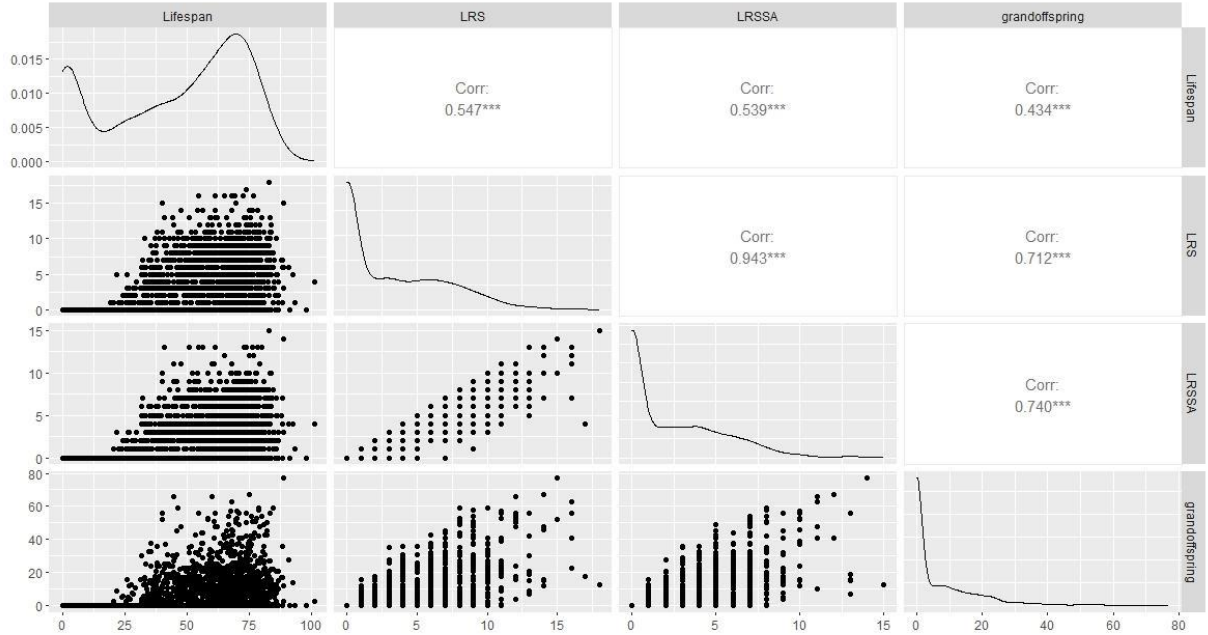

**Supplementary Figure 6: Pairwise correlations between fitness proxies (Lifespan, LRS, $LRS_{SA}$, and The number of grandoffspring) for 2790 individuals.** Pairwise Pearson correlation statistics (ρ) are shown above the diagonal with scatterplots with line of best fit (and SE indicated by the shaded area) shown below. Density plots showing distribution of fitness proxies are shown on the diagonal.

**Supplementary Table 4: Output from the beta model for mean offspring IGC with non-significant interactions included.** Displayed are the fixed and random effect estimates (posterior distribution median) with their 95% credible intervals (95% CrI); and the probability of Direction (*pd*) (not available for random effects). Significant effects (*pd* > 0.975) are shown in bold.

| Beta | | | |
|---|---|---|---|
| | Estimate | 95% CrI | pd |
| *Fixed effects* | | | |
| ***Intercept*** | **-9.032** | **[-9.318 - -8.728]** | **1.000** |
| ***LRS*** | **0.07** | **[0.053 - 0.088]** | **1.000** |
| ***Lifespan*** | **0.01** | **[0.006 - 0.013]** | **1.000** |

| | | | |
|---|---|---|---|
| ***Parish (Linthal)*** | **-0.217** | **[-0.423 - -0.009]** | **0.980** |
| *Sex (Male)* | -0.106 | [-0.249 - 0.04] | 0.926 |
| *Parish (Linthal) × Sex (Male)* | 0.09 | [-0.128 - 0.311] | 0.791 |
| *Random effects* | | | |
| *Parish-specific birth cohort (random intercept)* | 0.164 | [0.026 - 0.31] | |
| *Parish-specific birth cohort × LRS (random slope)* | 0.015 | [0.001 - 0.037] | |
